# Seed sequences mediate off-target activity in the CRISPR-interference (CRISPRi) system

**DOI:** 10.1101/2024.04.10.588881

**Authors:** Neha Rohatgi, JP Fortin, Ted Lau, Yi Ying, Yue Zhang, Mike Costa, Rohit Reja

## Abstract

The CRISPR interference (CRISPRi) system is a powerful tool that is widely used to selectively and efficiently silence genes in functional genomics research applications. However, the extent of off-target activity associated with the CRISPRi system has not been systematically investigated. Here, we utilized a genome-wide CRISPRi Cas9 single guide RNA (sgRNA) library to investigate the presence of off-target activity and its effects on gene expression. Our findings suggest that off-target effects in CRISPRi are quite pervasive and have direct and indirect impact on gene expression. Most of the identified off-target activities can be accounted for by complementarity of the PAM-proximal genomic sequence with the 3’ half of the sgRNA spacer sequence, the seed sequence. We also report that not all PAM proximal positions are equally important for binding of sgRNA to the DNA and that mismatches in the PAM-proximal regions can be tolerated depending on the sequence context.

## Introduction

Technology developments utilizing prokaryotic-derived Clustered Regularly Interspaced Short Palindromic Repeats (CRISPR) and the related Cas proteins have transformed the genome engineering and functional genomics landscapes. The widespread adaptability of this technology can be attributed to the fact that any genomic locus can be readily altered by using a combination of a nuclease protein and a single-guide RNA (sgRNA) to the target DNA ^1^. Since its initial report, CRISPR/Cas systems have rapidly evolved. They are not only limited to creating mutations and gene knock-outs through genome editing, but can also alter gene expression by modifying transcription ^2^. One such application is the development of CRISPR interference (CRISPRi) ^3^ wherein transcriptional repression of the target gene is achieved by fusing a repressive Krüppel-associated box (KRAB) domain to catalytically dead Cas9 nuclease (dCas9). By designing the sgRNAs to target the region near the transcription start site (TSS) of a gene, repression can be achieved by either blocking RNA Pol II elongation, or by blocking transcription initiation due to recruitment of co-repressor proteins by the KRAB domain ^3^. Typically, CRISPRi technology is employed in a high throughput screening format which involves the construction of a sgRNA library containing multiple guides that target promoter regions for each gene. This allows for a systematic evaluation of loss-of-function gene studies under a given perturbation. Additionally, CRISPRi has also been employed to repress single genes in induced pluripotent stem cells (iPSCs)^4^ and its potential across multiple gene therapy programs has been explored ^5^.

Although the CRISPR/Cas system and its dCas derivatives can effectively alter the function of the intended target, the presence of off-targets is still a major challenge. These off-targets occur as a result of unintended binding of sgRNAs to sequences that closely resemble the target sequence. Previous studies ^1,6^ employed chromatin immunoprecipitation sequencing (ChIP-seq) approaches to investigate the genome-wide binding of Cas9, and concluded that in some cases, sgRNA-directed Cas9 can bind to over 1000 sites in the genome. Still, not all sites undergo substantial cleavage as the nuclease activity of Cas9 requires an extensive base-pairing of the guide to the target DNA. Widespread binding of dCas9 might pose a substantial challenge for the CRISPRi system, as off-target binding, even if transient, could induce unintended transcriptional changes. The extent of off-target activity in the CRISPRi system has not been explored and no study has conclusively demonstrated a direct link between off-target sgRNA binding and its impact on gene expression. Additionally, it is reported that 10-12 nucleotides located at the 3’-end of sgRNA sequence, often referred as the “seed” sequence and presence of adjacent Protospacer Adjacent Motif (PAM) ^7^ sequence in the genome are crucial for target recognition ^8^. Whether all PAM proximal positions of the guide contribute equally to the target DNA recognition, and if this principle is true across all guides, still needs to be understood.

In this study, we utilized a genome-wide sgRNA library for the dCas9-KRAB CRISPRi system to interrogate the prevalence of off-target activity and its effect on gene expression. Our analysis confirmed the presence of off-targeting in the CRISPRi system and showed that it can have both direct and indirect impact on gene expression. Additionally, we found that not all PAM proximal positions are equally important for binding of sgRNA to the DNA and, depending on the sequence context, mismatches in the seed sequence of the guide do not necessarily impact its binding to the DNA.

## RESULTS

### Widespread off-targets in genome-wide CRISPRi screens

We applied genome-wide CRISPRi screening to identify new regulators of necrotic cell death pathways, particularly focusing on pyroptosis, an inflammatory cell death executed by the gasdermins ^9–11^, and necroptosis ^12,13^, an alternate form of inflammatory cell death mediated by RIPK1 (receptor interacting serine/threonine protein kinase 1), RIPK3, and its substrate pseudokinase, mixed lineage kinase domain-like (MLKL) ^14^. For the pyroptosis resistance CRISPRi screen, we infected an EA.hy926 cell line that had been engineered to express dCas9 fused with KRAB domain with the Horlbeck and Weissman *et al.* ^15^ whole-genome CRISPRi sgRNA library. After puromycin selection, cells were either electroporated with a control buffer or with LPS to induce cell death via pyroptosis (**Figure 1a**). We used an analogous procedure for the necroptosis resistance screen through which engineered Jurkat cells were infected with CRISPRi sgRNA library and treated with either DMSO or a cocktail of TNF, BV6, and Emricasan (TBE) to induce necroptosis (**Figure 1b**). The surviving cells from each screen were collected and sequenced individually, to identify the integrated lentiviral sgRNA sequence enriched in the surviving cell population. To detect top gene hits for a screen, most analysis tools aggregate the statistics across guides for a given gene and have the potential to miss the evidence of individual guide off-targeting. We addressed this problem, by performing our initial analysis only using the guide-level statistics (see Methods). The pyroptosis screen yielded expected top hits GSDMD and CASP4 which are both known regulators of this pathway ^16^ (**Figure 1c**). Similarly, in the necroptosis screen, MLKL, RIPK3, RIPK1, and TNFRSF1A emerged as the top hits, all of which are known players of this pathway ^17,18^ (**Figure 1d**). Upon examining other enriched guides in each screen, we noticed certain guides displayed a high fold change that deviated from the other guides designed to target the same gene (31 sgRNAs for pyroptosis resistance (**Figure 1c**) and 39 for necroptosis resistance (**Figure 1d**)). To better explain the unusual enrichment of these guides, we examined whether part of their sequence mapped to the promoters of genes that are known mediators of pyroptotic or necroptotic cell death. Using the 9-bp PAM proximal sequence of each guide, we performed a search for exact matches in the human genome (**Figure S1a**). Since CRISPRi sgRNAs are most effective when they target a region close to the TSS ^4^, we retained sgRNA matches that were within 2000 bp of a TSS. Among the most enriched guides in the pyroptosis screen, a total of 21 guides mapped to the promoter of GSDMD (plus an additional four guides with 8-bp seed sequences) (**Figure 1c**). In the necroptosis screen, 19 guides mapped to the promoter of RIPK3, 5 to MLKL, 2 to RIPK1, and 4 to TNFRSF1A promoter (**Figure 1d**). Using standardized statistical criteria to identity sgRNA hits across all genes targeted by the library (see Methods), we identified 76 potential off-targets in the pyroptosis screen, of which 35% (27 guides) mapped to the promoters of GSDMD and CASP4 (**Figure S1b**). Similarly for the necroptosis screen, 68% (35 out of 51 guides) of predicted off-targets mapped to the promoters of RIPK3, RIPK1, MLKL, or TNFRSF1A (**Figure S1c)**. To test whether a similar off-target effect is present in CRISPR/Cas9 knockout screens, we repeated the pyroptosis resistance screen using a CRISPR knockout (CRISPRn) library and EA.hy926 cells expressing wildtype Cas9; however, we did not see evidence of seed-sequence mediated off-targeting (**Figure S1d**). This suggested that off-target effects observed here might be exclusive to CRISPRi or similar dCas9 derivatives.

**Figure 1.**
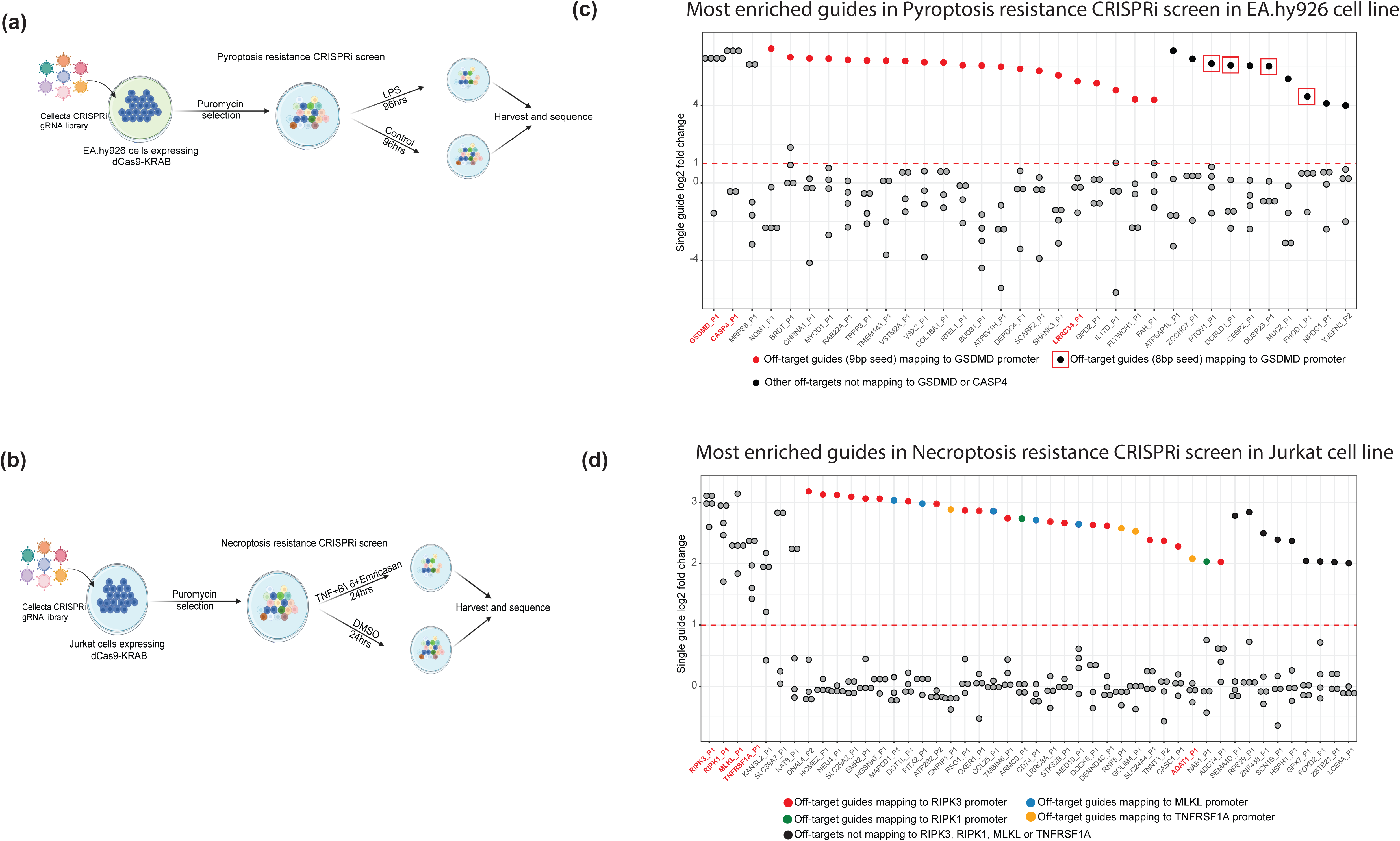
Pervasive CRISPRi off-target activity observed in pyroptosis and necroptosis resistance screens. (a) Schematics of pyroptosis resistance CRISPRi screen. (b) Same as (a) except for necroptosis resistance CRISPRi screen. (c) Maximally enriched guides from the pyroptosis resistance CRISPRi screen in EA.hy926 cell line. Each dot corresponds to a single guide. Guides whose 9 base-pair PAM proximal sequence exactly matches within a 2000 bp region centered around the GSDMD transcription start site (TSS) are colored red while the guides with 8bp seed matches are marked using a red square. All other off-targets are colored black. The x-axis is sorted by the following criteria: (a) multiple enriched guides that target the same gene are shown first, followed by guides whose 9bp PAM proximal region maps to GSDMD promoter and lastly all other enriched guides including the ones with 8bp PAM proximal matches to GSMD promoter. (d) Same as (c) except for the necroptosis CRISPRi screen in Jurkat cell line. Dots are colored red, blue, green, and orange if they map to the promoters of RIPK3, MLKL, RIPK1, and TNFRSF1A respectively. All other off-targets are colored black.

### PAM proximal 9 bp seed match alone is not sufficient for off-target activity

Our analysis indicated that the region of perfect complementarity between the promoter and the guide seed sequence can be as short as 8-9 bp to produce off-target effects. We next focused our attention on the sgRNAs that had perfect 9 bp PAM proximal seed sequence matches to the promoters of genes that mediate necrotic cell death. There were approximately 562 sgRNAs in the library with seed sequences that mapped to the GSDMD promoter (defined as -2000 to +2000 bps relative to the GSDMD TSS) in the pyroptosis screen, however, only 21 of these displayed fold-change values equivalent to that observed for the on-target sgRNAs (**Figure 2a**). This observation was also consistent for the necroptosis screen where only 4 out of 532 sgRNAs with seed sequence matches to the TNFRSF1A promoter displayed an effect equivalent to the on-target sgRNAs (**Figure 2b**). A similar trend was also observed at the promoters of the other known regulators of cell death (**Figure S2a-c**). Surprisingly, many of the sgRNAs that did not display a high fold change had seed sequence matches close to the TSS and in regions of open chromatin, suggesting that there must be factors in addition to 9-nt seed sequence complementarity that determine the binding stability of a sgRNA to genomic DNA. To investigate this further, we selected two guides from each screen: single guides targeting genes LRRC34 (sgLRRC34) and GRK4 (sgGRK4) from the pyroptosis screen and single guides targeting genes ADAT1 (sgADAT1) and HMGB1 (sgHMGB1) from the necroptosis screen.

**Figure 2.**
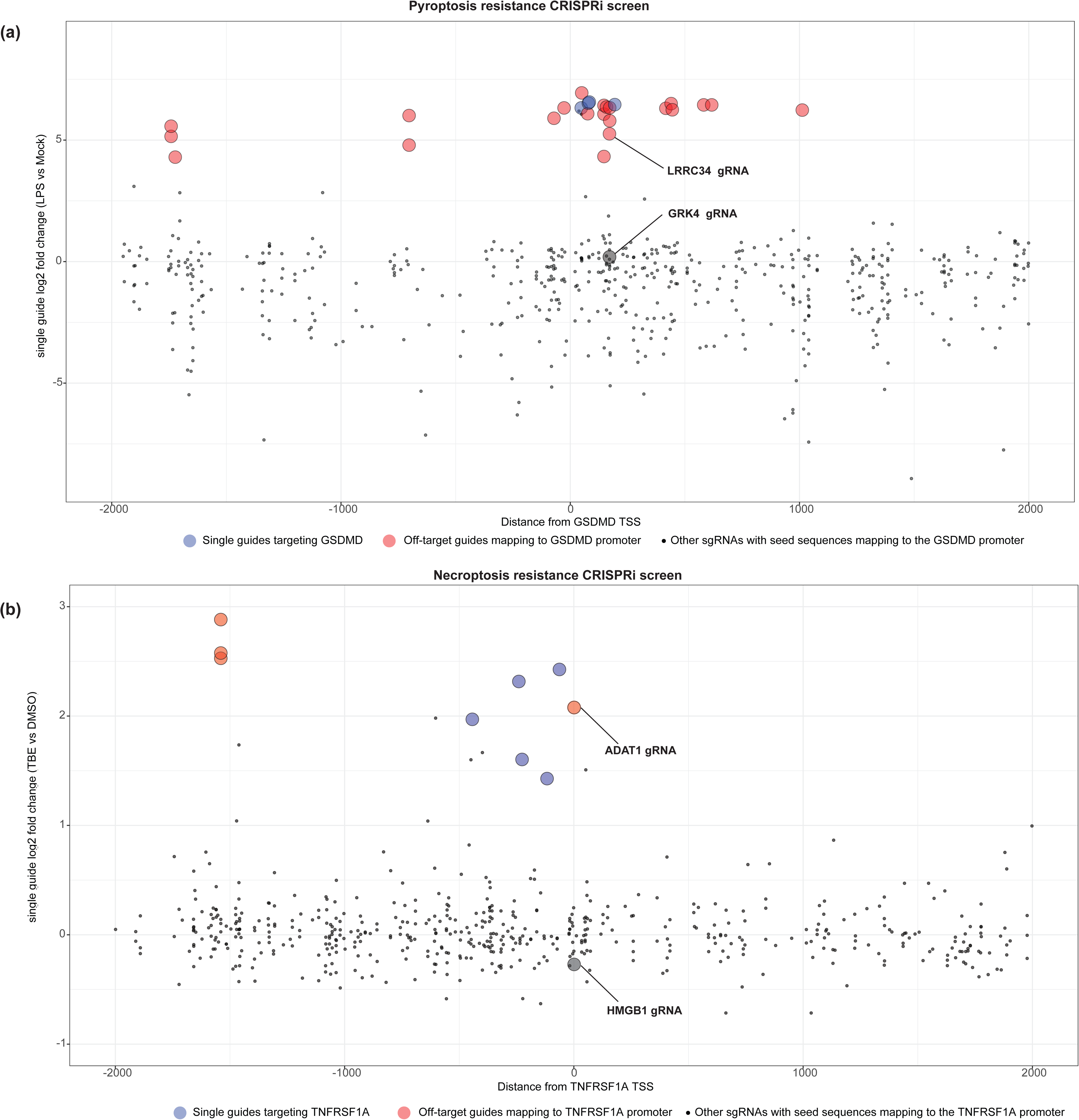
The off-target activity cannot be solely explained by the complementarity of the PAM-proximal 9 base-pair seed sequence to the promoter DNA. (a) Scatter plot with guides whose 9bp PAM proximal seed sequence exactly matches the region within a 2000 bp window centered around the GSDMD TSS. The x-axis shows the distance from the GSDMD TSS, while the y-axis shows the log2 fold change of individual guides from the pyroptosis resistance CRISPRi screen. Guides designed to target GSDMD are shown in blue, while off-target guides are colored in red. The guides sgLRRC34 and sgGRK4 were chosen for further investigation. (b) Same as (a) except for the necroptosis resistance CRISPRi screen. sgADAT1 and sgHMGB1 were identified for further investigation.

Guides sgLRRC34 and sgGRK4 were selected because they share an identical 9 bp seed match to the GSDMD promoter, however they have a 27-fold difference in their effect on cell survival following pyroptosis induction (log2 fold change for sgLRRC34 and sgGRK4 is 5.25 and 0.19 respectively, **Figure 2a**). Similarly, sgADAT1 and sgHMGB1 have a 7-fold difference in their effect on necroptosis survival, although they share the same seed match to the TNFRSF1A promoter (log2 fold change for sgADAT1 and sgHMGB1 is 2.07 and -0.27 respectively, **Figure 2b**). Based on these observations we suggest that exact 9 bp PAM-proximal complementarity, TSS proximity, and open chromatin might only partially determine sgRNA target recognition and off-target CRISPRi effects.

### ChIP-seq validates sequence-driven off-target activity of CRISPRi sgRNAs

With the aim of further characterizing the determinants of sgRNA binding, we designed a ChIP-seq experiment to identify genome-wide binding sites of dCas9 protein for each of the four single guides. dCas9-KRAB-expressing Jurkat cells were transduced with lentivirus containing either sgADAT1 or sgHMBG1. In parallel, dCas9-KRAB-expressing EA.hy926 cells were transduced with either sgLRRC34 or sgGRK4. dCas9 bound DNA fragments were immunoprecipitated and sequenced (**Figure 3a**). In the absence of any off-target activity, we anticipated seeing only one enrichment peak per guide, however, we observed that all four guides yielded multiple reproducible peaks ranging from 98 for sgGRK4 to 1141 for sgADAT1. Of these peaks, the proportion of promoter proximal peaks ranged from 10% for sgLRRC34 to 33% for sgGRK4 (**Figure 3b**). For each sgRNA, the strongest peak was detected at the intended target gene (**Figure S3a-d**). Only 48 peaks overlapped between the sgADAT1 and sgHMGB1 samples even though they had the same 9 bp seed sequence. There was only one overlapping off-target peak for sgLRRC34 and sgGRK4 (**Figure 3c**). Consistent with the off-target pyroptosis resistance effect of sgLRRC34, we observed a peak at the GSDMD promoter for sgLRRC34 but not for sgGRK4 (**Figure 3d)**.

**Figure 3.**
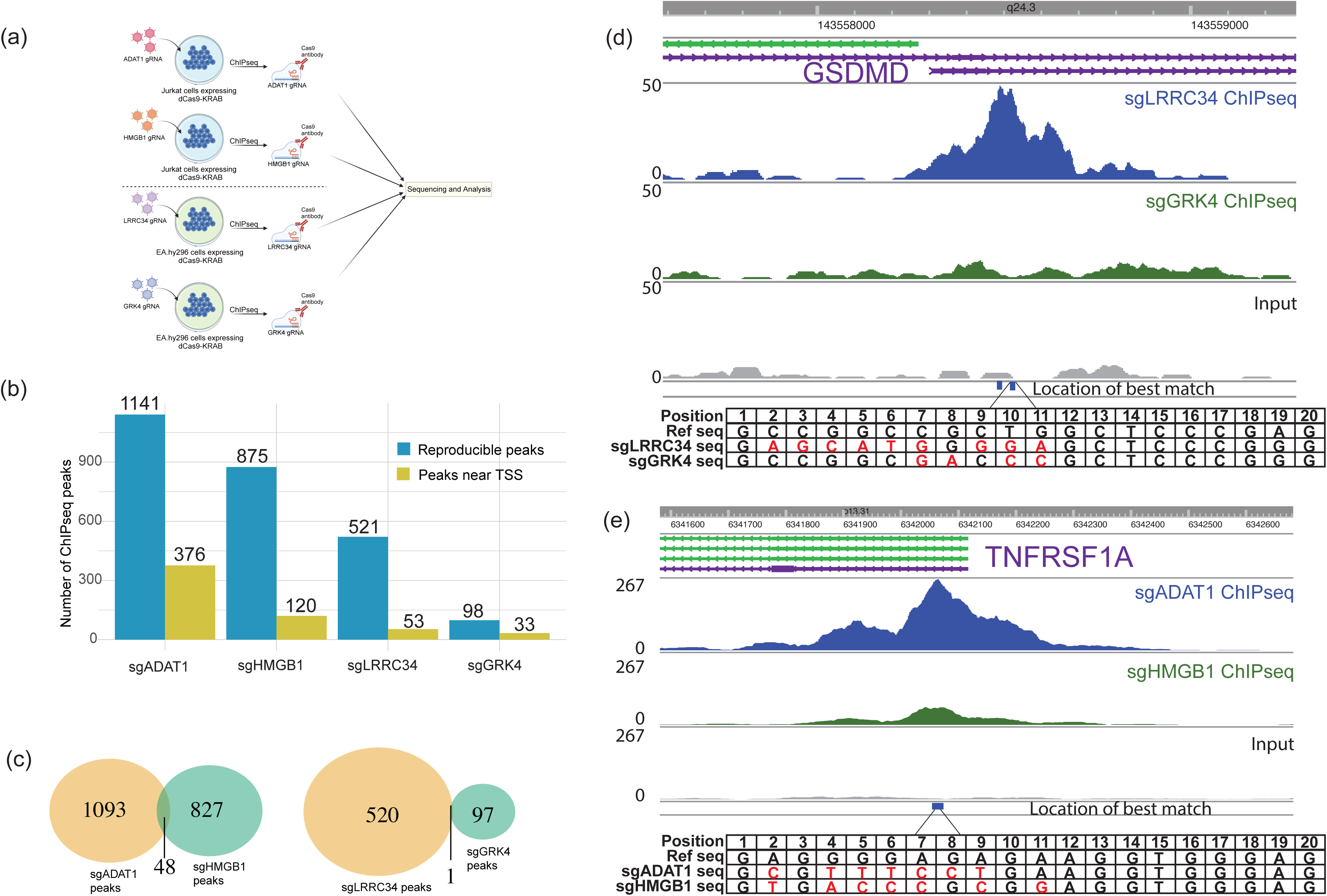
ChIP-seq validates the off-target activity observed in CRISPRi resistance screens. (a) Schematics of ChIP-seq experimental design for EA.hy926 and Jurkat cells. (b) Barplot comparing the number of reproducible ChIP-seq peaks observed for each single guide experiment (in blue) and peaks within a 2000 bp window of any TSS (in yellow). (c) Left panel: Venn diagram showing ChIP-seq peak overlap between sgADAT1 (in orange) and sgHMGB1 (in green); Right panel: Same as left panel, except for sgLRRC34 and sgGRK4. (d) Graph showing ChIP-seq reads at the GSDMD promoter for sgLRRC34 (blue), sgGRK4 (green), and Input (grey). Bottom panel: The sequences of sgLRRC34 and sgGRK4 are evaluated in comparison to the sequence found at the most optimal match location within the GSDMD promoter. (e) Same as (d) except at the TNFRSF1A promoter for sgADAT1 (blue), sgHMGB1 (green), and Input (grey). Bottom panel: The sequences of sgADAT1 and sgHMGB1 are evaluated in comparison to the sequence found at the most optimal match location within the TNFRSF1A promoter.

While both sgRNAs share the same 9 bp seed sequence, sgGRK4 actually has more matches to the genomic protospacer sequence 5’ to the seed region (**Figure 3d**, lower panel). However, there is an additional genomic sequence just a few bp upstream, that has a single mismatch at position 19 of the seed sequence (**Figure 3d).** Importantly, this site has complementarity with sgLRRC34, but not sgGRK4, that extends an additional 3 bp 5’ to the 9-bp seed sequence (**Figure 3d, S3e**). The ChIP-seq data does not have sufficient resolution to distinguish the dCas9 binding occupancy between these two sites, and the binding motif analysis described below suggests that both sites may be bound by sgLRRC34. We found sgADAT1- and sgHMGB1-mediated dCas9 peaks at the TNFRSF1A promoter, however the sgADAT1 peak was much greater in intensity, possibly explained by extension of the complementarity by 2 bp 5’ to the 9 bp seed sequence (**Figure 3e**, lower panel). Extensive base-pairing between the sgRNA and the target DNA beyond the 9 bp seed sequence is required for substantial DNA cleavage by wildtype Cas9 in the CRISPR knockout system ^1^, but it remains to be defined how this additional complementarity influences occupancy of the dCas9 complex on genomic DNA.

### ChIP-seq integration with RNA-seq confirms widespread off-target activity of the CRISPRi system

To assess the direct and indirect effects of dCas9-sgRNA off-target binding on gene expression, we performed a differential RNA-seq analysis to compare the individual sgRNA transduced samples to the uninfected samples for each of the four guides in our study. Although the CRISPRi sgRNA library was designed to minimize off-target effects driven by 1-2 of 19 bp mismatches, we discovered that the number of differentially expressed genes varied greatly, from as few as 88 genes for cells transduced with sgADAT1, to roughly 2088 genes for cells transduced with sgLRRC34 (FDR < 0.05; **Figure S4a**). This broad impact on gene expression could potentially be due to not only the direct effect of sgRNAs binding near the promoters of genes and repressing them, but also the indirect effect of the repressed genes which in turn affect the expression of other genes. To estimate the direct effects, we assigned each single guide ChIP-seq peak to its closest gene and looked at the impact of this binding on the proximal gene’s expression. As expected, a strong sgADAT1 peak at the ADAT1 gene promoter, and a weaker peak at the TNFRSF1A promoter, correlated with a substantial reduction in ADAT1 transcript, as well as robust suppression of TNFRSF1A transcript (**Figure 4a**). Additionally, five other genes appear to be impacted by off-target binding of sgADAT1 to their promoters. This number was higher for sgLRRC34, where the repression of 20 genes (out of 2088 differentially expressed genes) appears to be due to direct effects of off-target binding (**Figure 4b**). Although a sgLRRC34 peak was observed at the LRRC34 gene promoter, it is important to note that no gene suppression was observed since LRRC34 is not expressed in the EA.hy926 cell line (**Figure S3c and 4b**). 14 (out of 297) and 23 (out of 938) differentially expressed genes can be attributed to the direct off-target binding of sgGRK4 and sgHMGB1, respectively (**Figure S4b,c**). While the numbers of direct targets impacted by the off-targeting activity are relatively modest, RNA-seq data indicates that these could lead to indirect effects on many more genes. Collectively, these findings suggest that pervasive off-target activity in CRISPRi can result in extensive alterations in the transcriptome.

**Figure 4.**
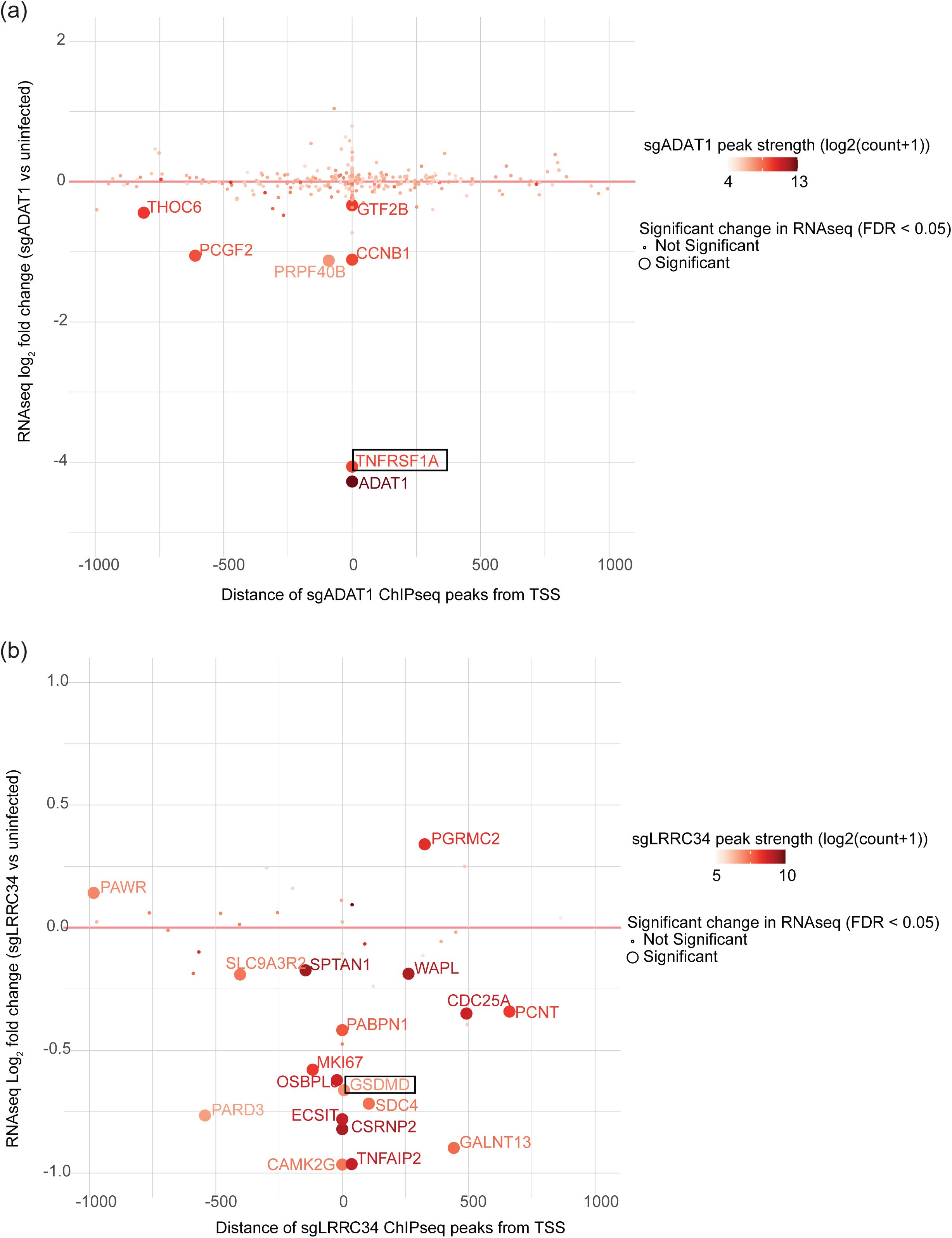
Integration of ChIP-seq and RNA-seq shows direct effect of off-target binding. (a) Scatter plot showing the distribution of sgADAT1 ChIP-seq peaks relative to the TSS (on the x-axis) and log2 fold change values from RNA-seq (on the y-axis) for the closest genes near the sgADAT1 ChIP-seq peaks. Dots are colored by the strength of ChIP-seq peak signal and are bigger in size if found significantly changed in RNA-seq (FDR < 0.05). (b) Same as (a) except for sgLRRC34 ChIP-seq peaks.

### Variable contribution of seed sequence nucleotide positions to DNA binding

To determine if all PAM-proximal positions of the sgRNA are equally important for DNA binding, we performed a *de novo* motif search for each single guide ChIP-seq peak dataset using the MEME suite. Interestingly, MEME motifs for all four guides captured variable lengths of the PAM-proximal region of their respective sgRNA sequences (**Figure 5a,b and S5a,b**). Consistent with the previous literature ^7^, the 3’ genomic NGG PAM sequence was present in the MEME motifs for all four guides. For sgADAT1, only nucleotides at positions 16-20 in the sgRNA sequence were strongly enriched in the target DNA, suggesting that these positions might be critical for binding. (**Figure 5a**). Additionally, nucleotides at PAM-proximal positions 10-15 may contribute to DNA binding, albeit with more tolerance for mismatches. In contrast, for sgHMGB1 containing the same 9 bp seed sequence as sgADAT1, the nucleotides at PAM proximal positions 10-20 in the sgRNA sequence appear to be important for DNA binding and were less tolerant of mismatches (**Figure 5b**). For both sgRNAs, the strength of dCas9 peaks diminished as more mismatches were observed in the PAM proximal region (**Figure 5c,d)**. For sgLRRC34 and sgGRK4, the PAM proximal positions 9-20 in the sgRNA were enriched in the target DNA, however, mismatches at position 18 did not robustly diminish the peak strength (**Figure S5a,b,c,d**). sgLRRC34 and sgGRK4 have a higher GC content in the 9-nt seed sequence than sgADAT1 and sgHMGB1 (1 vs. 3 A/T nucleotides) which could create a more stable DNA-RNA hybrid that can better tolerate this mismatch, but it is not clear why mismatches as position 18 would be preferentially tolerated. Together, this data suggests that the stability of off-target binding is driven by the PAM-proximal seed sequence positions, but that the length of this seeding sequence and the tolerance of mismatches at different positions varies by sgRNA. To more precisely define the features, including GC content, that control this variation, a more diverse set of sgRNAs will need to be analyzed for binding and CRISPRi activity.

**Figure 5.**
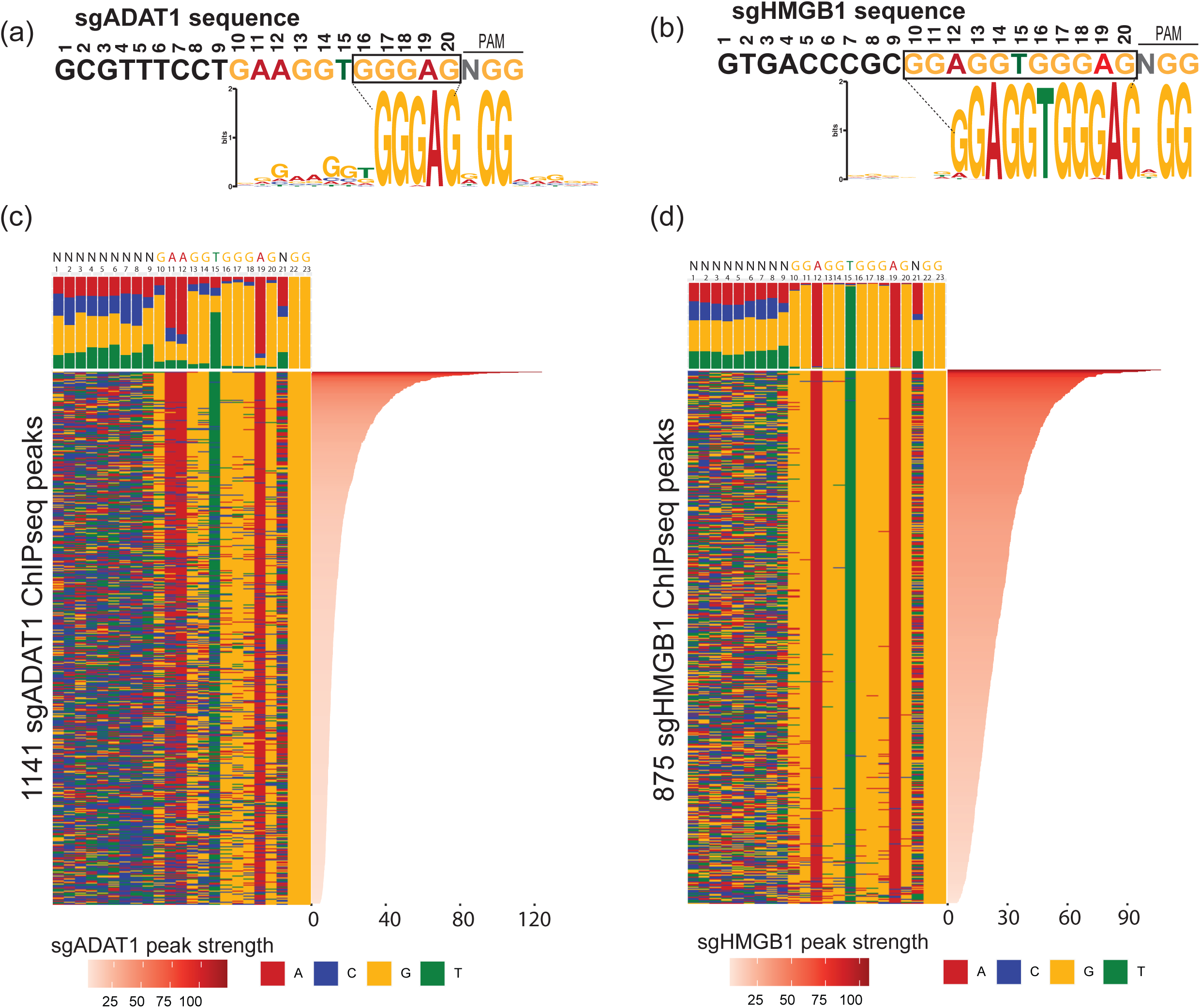
Sequence analysis of ChIP-seq peaks reveals genomic DNA binding specificity with variable contribution of sgRNA nucleotide positions. (a) Sequence logo showing the top MEME motif enriched in the sgADAT1 ChIP-seq peaks. For reference, sgADAT1 sequence is shown at the top and colored to reflect an enrichment of nucleotides observed in the sequence motif. Box highlights the nucleotides most critical for target recognition. (b) Same as (a) except for sgHMGB1. (c) Color chart representation of 23 bp of DNA sequences enriched in all 1141 sgADAT1 ChIP-seq peaks, sorted by the peak strength (shown in the right panel). The frequency of nucleotides A (red), T (green), G (orange), and C (blue) at each position in the gRNA are shown in the top panel. (d) Same as (c) except for sgHMGB1.

## Discussion

In this paper, we show that the off-target activity of the CRISPRi system is more prevalent than previously recognized, in part because dCas9 can exert off-target activity driven by seed sequence complementarity analogous to siRNA ^19^ and distinct from CRISPRko (wildtype Cas9). Some of the off-targets detected in our screens could be readily explained by identifying the guide’s 9 bp seed sequence match to the promoter region of a top-ranking on-target gene hit. However, other sgRNAs in the library with identical 9 bp seed sequences did not confer off-target cell survival phenotypes. This indicates that seed sequence complementarity is only part of what determines stable sgRNA target recognition.

Using ChIP-seq we showed that sequence complementarity that extends as far as 12 bp of seed sequence can sometimes be crucial for a strong occupancy of dCas9 on the DNA and for off-target transcriptional repression. By integrating ChIP-seq and RNA-seq data together, we demonstrated that although the direct repressive effects of off-target binding may be limited, they can lead to more extensive secondary changes in the transcriptome. Using the dCas9-bound enriched sequences for each of the single guides, we showed that not all PAM proximal positions are equally important for target DNA binding, and that mismatches can be tolerated in the seed region, possibly depending on the sequence composition of the PAM proximal region.

Given that the off-target activity largely depends on the sgRNA seed sequence, we hypothesize that other dCas9 derivatives, including the CRISPR activation (CRISPRa) system, may be susceptible to similar effects. Caution should be exhibited when looking at top hits from a CRISPRi high throughput screen as off-targets with high enrichment scores may bias the overall ranking of genes. Additionally, CRISPRi/a sgRNAs could lead to false interpretations of gene perturbation phenotypes when employed to repress individual genes specially in the context of therapeutics.

Several approaches can be used to minimize off-target activity in the context of individual gene perturbations, such as the use of dual guides ^8^, engineering dCas9 to increase specificity ^20^, or by controlling experimental conditions like the concentration of dCas9 or the sgRNA ^21^. In addition, computational approaches can also be utilized to predict off-target binding sites of a sgRNAs, by searching for promoter matches to the 9-12 bp seed sequence region. Specifically, for CRISPRi/a, we can leverage this information to avoid using sgRNAs with high predicted off-target activity. In conclusion, any transcriptomics or phenotypic data derived from dCas9 perturbation technologies should be interpreted with a careful understanding of these seed-sequence-mediated off-target effects.

## Materials and Methods

### Pyroptosis resistance CRISPRi screen

The human epithelial EA.hy926 CRISPRi cell line was engineered to constitutively express dCas9-KRAB and infected with Cellecta (Mountain View, CA, USA) CRISPRi human genome-wide library at MOI of 0.33-0.44 and a representation of 200-250 cells per sgRNA for each of 3 replicates. The cells were passaged for 16 days, selected with 0.5ug/ml puromycin, followed by electroporation with either LPS (2.0 x 10^8^ cells per replicate) or control buffer (4.5 x 10^7^ cells per replicate). Electroporation was performed according to Lee et al., 2018 ^22^, with 1ug LPS per 1 x 10^6^ cells in aliquots of 7 x 10^6^ cells per electroporation. LPS electroporation resulted in pyroptotic cell death of 99.3-99.6% of cells. Cells were harvested 3 days after infection (reference samples) and 4 days after treatments. For all cell collections, genomic DNA isolation, PCR amplification of the sgRNA sequences, amplicon purification, and next-generation paired-end sequencing were performed according to Callow *et al*., 2018 ^23^. Sufficient genomic DNA was used for PCR amplification to maintain a representation of approximately 200 cells per sgRNA for reference and control electroporation samples (with the exception of one reference replicate at 100 cells per sgRNA).

### Necroptosis resistance CRISPRi screen

Human Jurkat CRISPRi cell line, which had been engineered to express dCas9-KRAB constitutively, were infected with Cellecta (Mountain View, CA, USA) CRISPRi human genome-wide library at MOI=0.6 with a target of 350-fold gRNA representation. Cells were selected by puromycin for three days and recovered for another two days. On post-infection day 7, cells were treated in triplicate with a combination of necroptotic stimuli TBE (100 ng/ml human TNF, 1 uM BV6 and 10 uM Emricasan) to induce cell death by necroptosis or with DMSO for 24 hrs. Cells were collected in triplicate via the following method: a) reference sample before puromycin selection, b) another reference collected right before TBE treatment, c) DMSO control, collected 24 hours post-treatment, and d) TBE treatment samples, collected 24 hrs post-treatment. Following cell harvest, genomic DNA was isolated and PCR amplification of integrated gRNAs was performed.

### CRISPRi screen data analysis

CRISPRi screen data was analyzed using the crisperVerse R package ^24^. For the pyroptosis screen, guides with log2 fold change > 3.5 and FDR < 0.05 in LPS treated samples relative to mock and with log2 fold change > 2 and FDR < 0.05 relative to the reference samples were considered as highly enriched. Similarly, for the necroptosis screen, guides with log2 fold change > 2 and FDR < 0.05 in TBE treated samples relative to DMSO and with log2 fold change > 1 and FDR < 0.05 relative to the reference samples were considered highly enriched. The log2 fold change thresholds were chosen to find the maximum number of off-targets (*i.e*., single guide RNA per gene hits). The 9 bp PAM proximal seed sequences for the most enriched guides were searched against the whole human genome using the crisprBowtie R package ^24^. We assigned the most enriched guides to genes if the 9 bp PAM proximal sequence of the guide matched to the promoters (defined as +/- 2000 bp centered around the TSS) of known mediators such as GSDMD and CASP4 for pyroptosis, and RIPK3, RIPK1, MLKL, and TNFRSF1A for the necroptosis.

To comprehensively look for off-targets across the whole CRISPRi library, we employed the following criteria for each screen. A guide was designated as having an off-target if it’s log2 fold change was outside the range defined as: Q1 – 1.5*IQR to Q3 + 1.5*IQR, where Q1 and Q3 represent first and third quartiles of the log2 fold change for all sgRNAs targeting the gene, and IQR is defined as Q3 – Q1. The IQR was computed using the log2 fold change values for all guides targeting the same genes. Additionally, we also required that the guide designated as off-target had a log2 fold change > 2.

### ChIP-seq protocol

Spacer sequences GAGCATGGGGAGCTCCCGGG (sgLRRC34) and GCCGGCGACCCGCTCCCGGG (sgGRK4) were cloned into vector SHC201 and packaged into lentivirus by Sigma-Aldrich (Saint Louis, MO, USA). EA.hy926 dCas9-KRAB cells were transduced in duplicate samples with each lentivirus and selected with 0.45 ug/ml puromycin starting 3 days after transduction. Control uninfected cells were similarly processed in parallel, but without puromycin selection. Each cell sample was expanded to 3 x 10^7^ cells and fixed in flasks in growth medium with 1.1% formaldehyde, 0.01 M NaCl, 0.1 mM EDTA, 5 mM HEPES, pH 7.9 for 15 min. Cells were then incubated in 0.125 M glycine for 5 min, scraped off the flasks with a rubber policeman, and kept on ice for the remainder of the protocol. Cells were centrifuged at 800 x g for 10 min, resuspended in PBS, 0.5% Igepal, 0.1 mM PMSF. Cells were centrifuged again and frozen on dry ice after removing supernatant. An analogous procedure was implemented for Jurkat dCas9-KRAB cells with spacer sequences sgADAT1 (GCGTTTCCTGAAGGTGGGAG) or sgHMGB1 (GTGACCCGCGGAGGTGGGAG).

Samples were sent to Active Motif (Carlsbad, CA, USA) for ChIP-Seq. Active Motif prepared chromatin, performed ChIP reactions, generated libraries, sequenced the libraries and performed basic data analysis. In brief, cells were fixed with 1% formaldehyde for 15 min and quenched with 0.125 M glycine. Chromatin was isolated by adding a lysis buffer, followed by disruption with a Dounce homogenizer. Lysates were sonicated and the DNA sheared to an average length of 300-500 bp with Active Motif’s EpiShear probe sonicator (cat# 53051). Genomic DNA (Input) was prepared by treating aliquots of chromatin with RNase, proteinase K and heat for de-crosslinking, followed by SPRI beads clean up (Beckman Coulter) and quantitation by Clariostar (BMG Labtech). Extrapolation to the original chromatin quantity allowed determination of the total chromatin yield.

An aliquot of chromatin (30 ug) was precleared with protein G agarose beads (Invitrogen). Genomic DNA regions of interest were isolated using 4 ug of antibody against dCas9 protein (Active Motif 61757, lot 10216001). Complexes were washed, eluted from the beads with an SDS buffer, and subjected to RNase and proteinase K treatment. Crosslinks were reversed by incubation overnight at 65°C, and ChIP DNA was purified by phenol-chloroform extraction and ethanol precipitation. Quantitative PCR (qPCR) reactions were carried out in triplicate on specific genomic regions using SYBR Green Supermix (Bio-Rad). The resulting signals were normalized for primer efficiency by carrying out qPCR for each primer pair using Input DNA. Illumina sequencing libraries were prepared from the ChIP and Input DNAs on an automated system (Apollo 342, Wafergen Biosystems/Takara). After a final PCR amplification step, the resulting DNA libraries were quantified and sequenced on Illumina’s NextSeq 500 (75 nucleotide reads, single end).

### ChIP-seq data analysis

Raw FASTQ files were aligned to the human reference genome (GRCh38) using BWA (version 0.7.12-r1039) ^25^. Reads that mapped uniquely to the genome and aligned with no more than 2 mismatches were used for downstream analysis. MACS2 (version 2.1.0) ^26^ was used to call the peaks relative to the pooled input. Motif analysis was performed using the MEME suite tool ^27^. WashU epigenome browser ^28^ was used to look at locus specific ChIP-seq read enrichment. R package ChIPpeakAnno ^29^ was used to compute the significance of overlap between ChIP-seq peaks. Peaks were assigned to the closest gene if they were within +/-1000 bp of a known TSS using BEDTools^30^.

### Identifying the best sgRNA match in the ChIP-seq peaks

The best match for a guide RNA to the genomic DNA within each ChIP-seq peak was estimated using a weighted score of matches at each position in the 20 bp sgRNA sequence. The score adds more weight to the matches closer to the PAM proximal region than matches in the PAM distal region. It is calculated as the sum of all the matched positions multiplied by the total number of matches of the guide sequence with the genomic reference.

Weighted Score = SUM(match position) * total number of matches

### RNA-seq protocol

EA.hy926 dCas9-KRAB cells infected with lentivirus carrying either the sgLRRC34 or sgGRK4 as described above were plated into a 6-well plate, 4 days after starting puromycin selection, and RNA was extracted with RNeasy Plus Mini Kit (74134, QIAGEN) according to the manufacturer’s instructions once cells reached approximately 90% confluence. The experiments were performed in duplicates and uninfected samples were used as controls. A similar process was implemented for Jurkat cells expressing a dCas9-KRAB fusion protein with infection performed using lentivirus containing either sgADAT1 or sgHMGB1 and in triplicates. Total RNA was isolated from all the samples and quantified with a Qubit RNA HS Assay Kit (Thermo Fisher Scientific, Waltham, MA, USA) and quality was assessed using RNA ScreenTape on TapeStation 4200 (Agilent). For sequencing library generation, the Truseq Stranded mRNA kit (Illumina, San Diego, CA, USA) was used with an input of 100-1000 nanograms of total RNA. Libraries were quantified with a Qubit dsDNA HS Assay Kit (Thermo Fisher Scientific, Waltham, MA, USA) and the average library size was determined using D1000 ScreenTape on TapeStation 4200 (Agilent Technologies). Libraries were pooled and sequenced on NovaSeq 6000 (Illumina) to generate 30 million single-end 50-bp reads for each sample.

### RNA-seq data analysis

GSNAP ^31^ (version 2013-11-10) was used to align raw FASTQ reads to the human reference genome (GRCh38/hg38) with the following parameters “-M 2 -n 10 -B 2 -i 1 -N 1 -w 200000 -E 1 --pairmax-rna=200000 --clip-overlap”. Reads were filtered to include only the uniquely mapped reads. Limma ^32^ R package was used to perform differential expression analysis.

## Acknowledgements

We thank Vishva Dixit, Kim Newton and Dorothee Nickles for discussion and feedback. This work was funded by Genentech/Roche.

## Declaration of interests

NR, JP, TL, MC and RR, are current employees of Roche/Genentech. YZ is an employee of Scribe therapeutics and YY is an employee of Sana Biotechnology.

## Data Availability

RNA-seq and ChIP-seq data is available in GEO under the accession numbers: GSE252977 and GSE252978.

---

**Figure S1.**
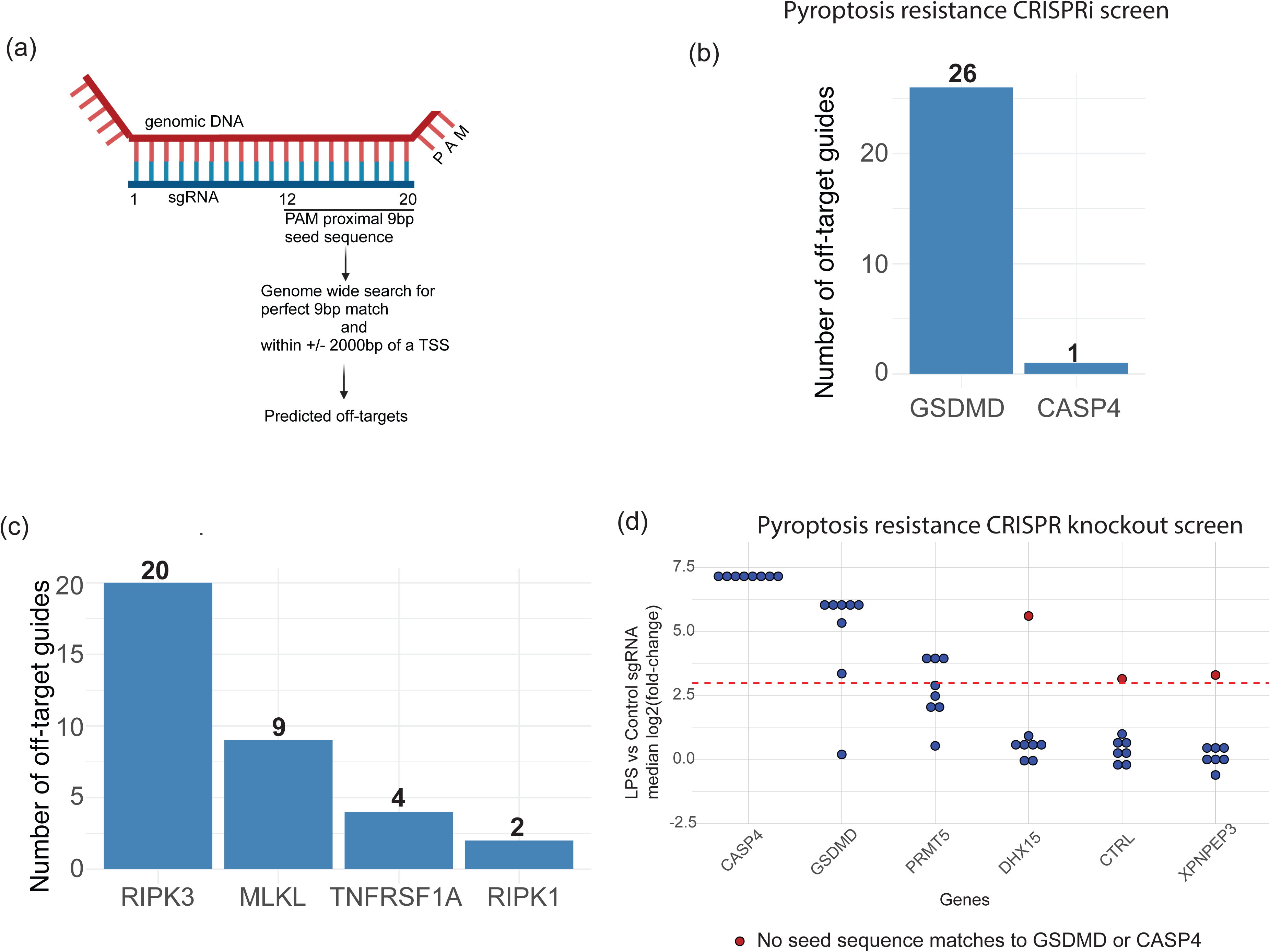
Assessment of off-target activity in the whole-genome CRISPRi library, related to Figure 1. (a) Schematics depicting search criteria used to predict off-targets. (b) Barplot showing the number of enriched gRNAs (on the y-axis) from the whole library with seed sequences that map to the promoters of GSDMD and CASP4 (on the x-axis) in the pyroptosis resistance CRISPRi screen. (c) Same as (b) except for the necroptosis resistance CRISPRi screen. (d) Graph showing the log2 of fold change for all guides of the six genes that showed at least one sgRNA with log2(fold change) > 3 for LPS vs control treatment in the pyroptosis resistance CRISPR knockout (wildtype Cas9) screen.

**Figure S2.**
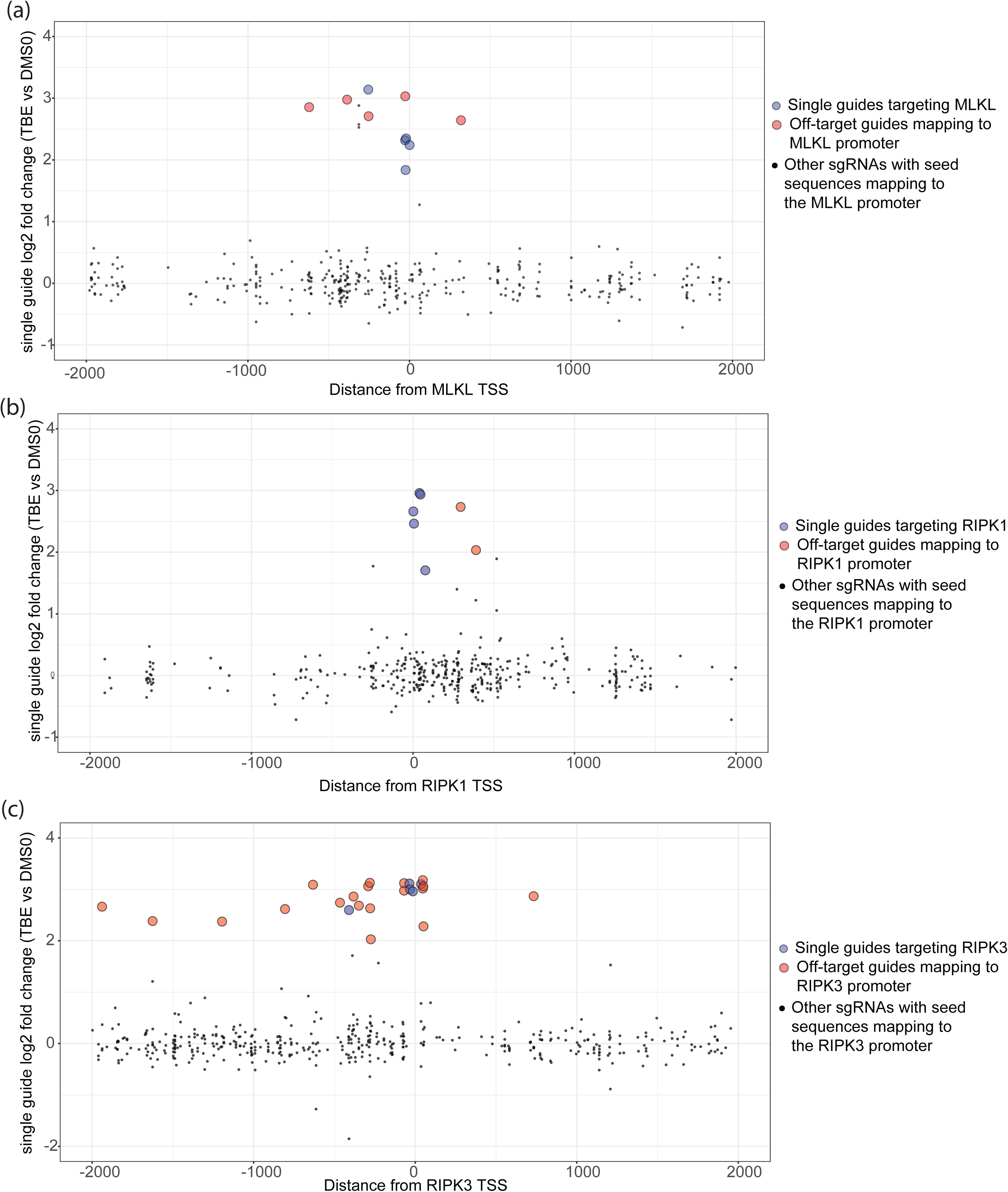
CRISPRi on and off-targets at the promoters of other top hits from both screens, related to Figure 2. (a) Scatter plot with guides whose 9bp PAM proximal seed sequence exactly matches the region within a 2000 bp window centered around the MLKL TSS. The x-axis indicates the distance from MLKL TSS, while the y-axis shows the log2 fold change values for individual guides from the necroptosis resistance CRISPRi screen. Guides designed to target MLKL are shown in blue, while off-target guides are colored in red. (b) Same as (a) except for RIPK1. (c) Same as (a) except for RIPK3.

**Figure S3.**
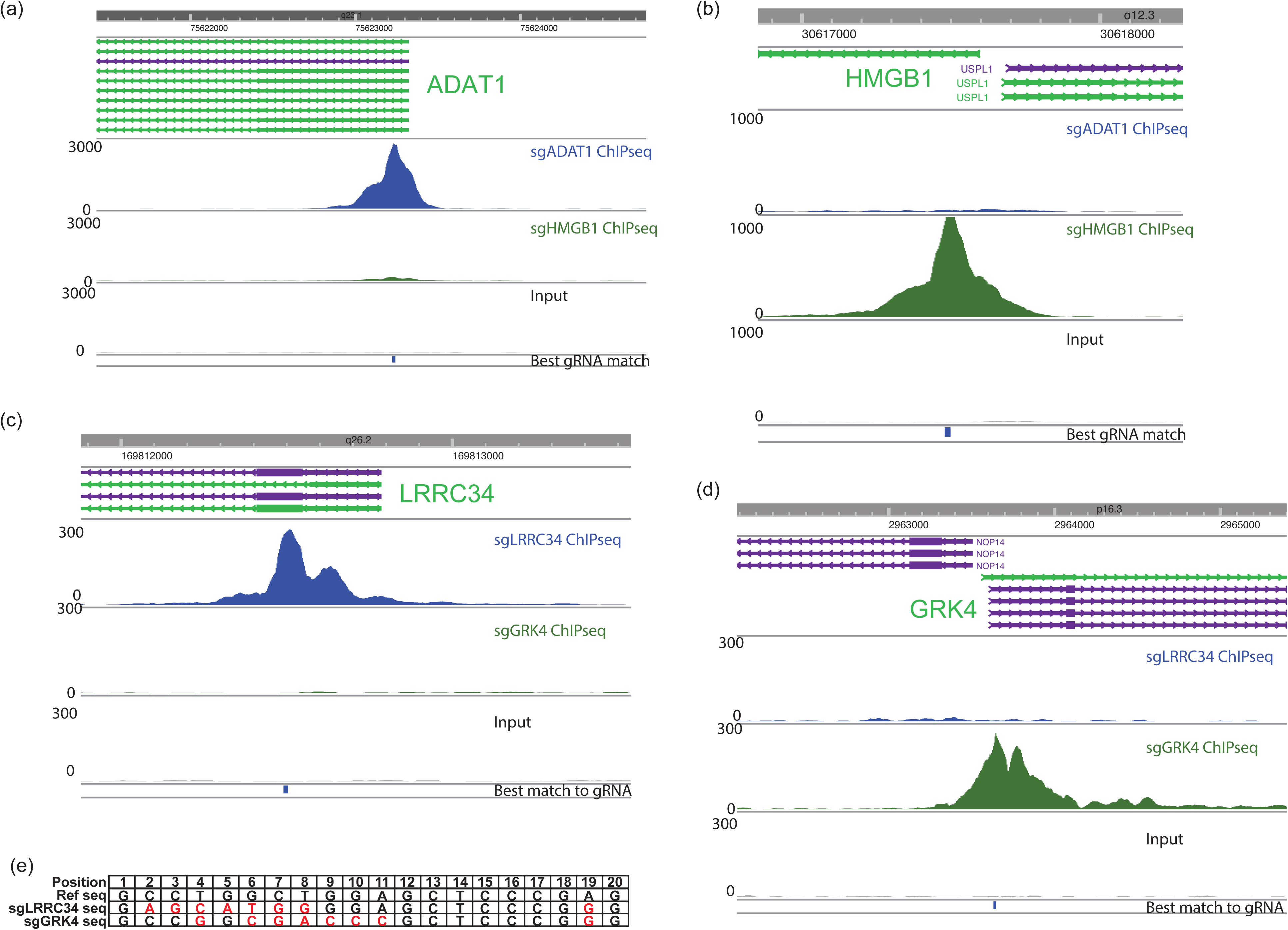
ChIP-seq peak occupancy at the intended on-target locus, related to Figure 3. (a) Graph showing ChIP-seq reads at the ADAT1 promoter for sgADAT1 (blue), sgHMGB1 (green), and Input (grey). (b) Same as (a) except at HMGB1 promoter. (c) Graph showing ChIP-seq reads at the LRRC34 promoter for sgLRRC34 (blue), sgGRK4 (green), and Input (grey). (d) Same as (c) except at GRK4 promoter. (e) The second sequence match of sgLRRC34 and sgGRK4 are evaluated in comparison to the genomic sequence found within the GSDMD promoter.

**Figure S4.**
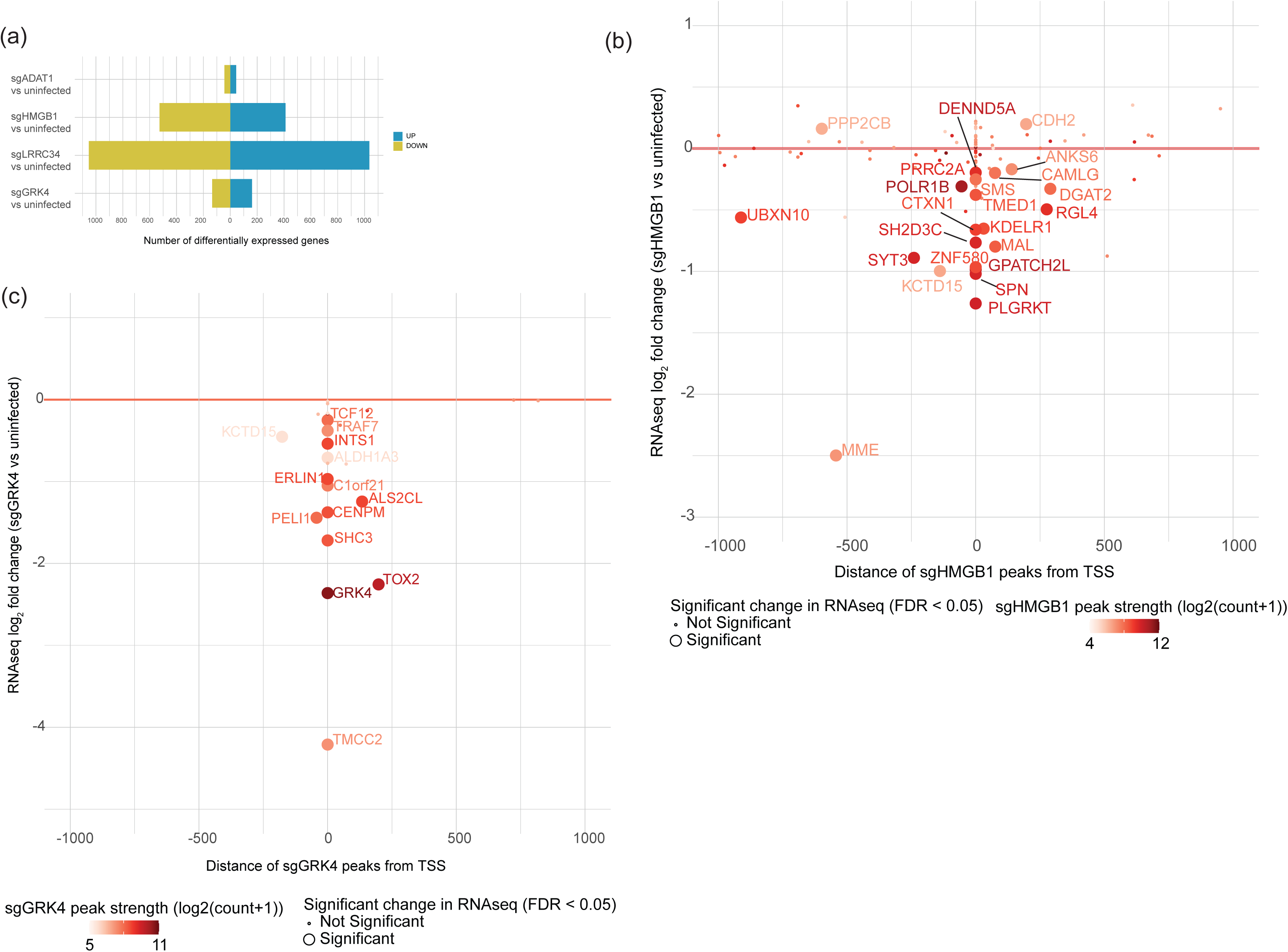
RNA-seq confirms the suppressive effect of off-target ChIP-seq peaks, related to Figure 4. (a) Bar plot showing the number of differentially expressed genes (FDR < 0.05) for sgADAT1, sgHMGB1, sgLRRC34, and sgGRK4 compared to uninfected samples. Up-regulated genes are shown in blue and down-regulated genes are shown in yellow. (b) Scatter plot showing the distribution of sgHMGB1 ChIP-seq peaks relative to the TSS (on the x-axis) and log2 fold change values from RNA-seq (on the y-axis) for the closest genes near the sgHMGB1 ChIP-seq peaks. Dots are colored by the strength of ChIP-seq peak signal and have large size if found significantly changed in RNA-seq (FDR < 0.05). (c) Same as (b) except for sgGRK4.

**Figure S5.**
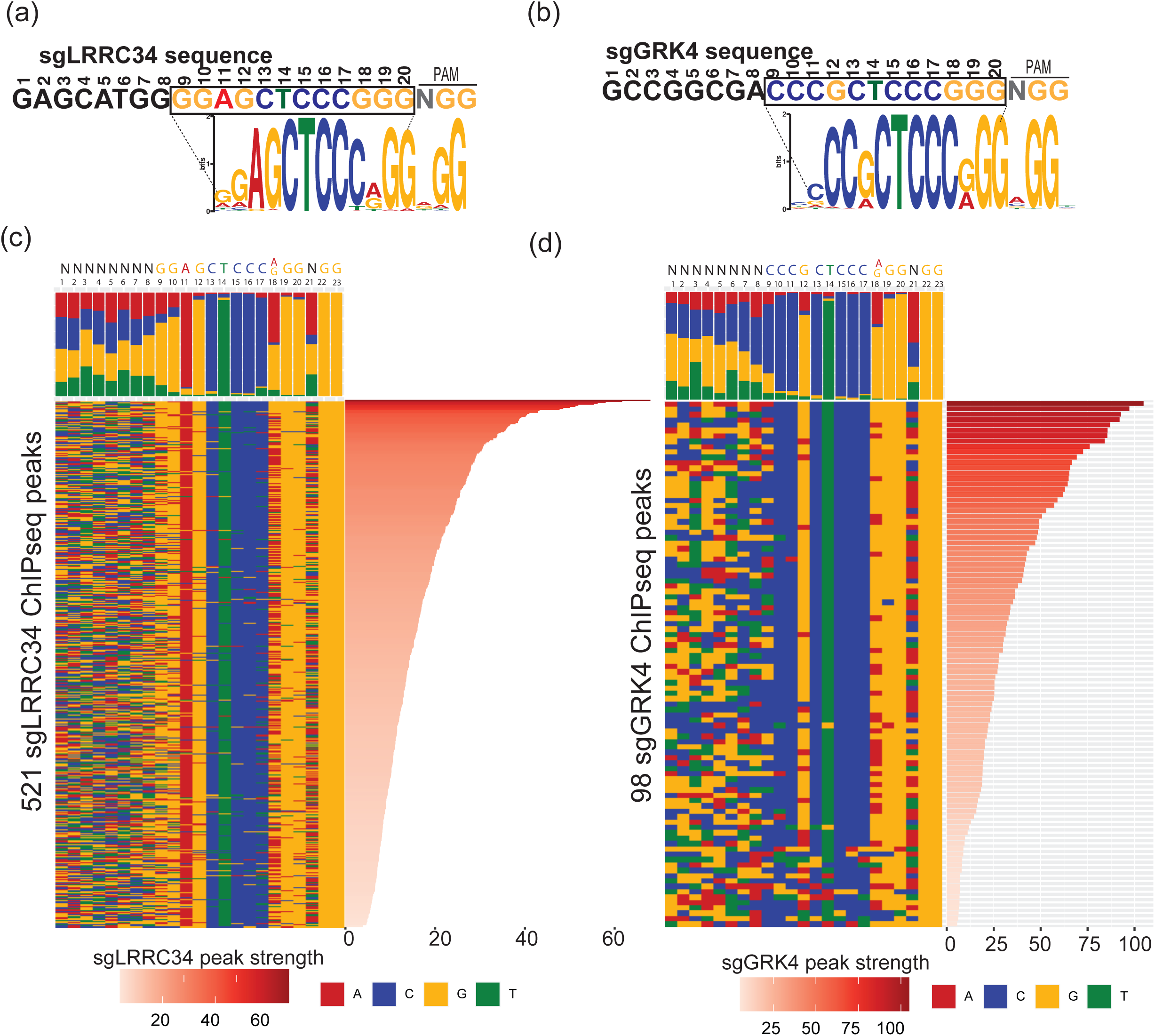
Sequence analysis of dCas9-bound single guide peaks reveals variable nucleotide specificity, related to Figure 5. (a) Sequence logo showing the top MEME motif enriched in the sgLRRC34 ChIP-seq peaks. For reference, sgLRRC34 sequence is shown at the top and colored to reflect an enrichment of nucleotides observed in the sequence motif. Box highlights the nucleotides most critical for target recognition. (b) Same as (a) except for sgGRK4. (c) Color chart representation of 23 bp of DNA sequences enriched in all 521 sgLRRC34 ChIP-seq peaks, sorted by the peak strength (shown in the right panel). The frequency of nucleotides A (red), T (green), G (orange), and C (blue) at each position in the gRNA are shown in the top panel. (d) Same as (c) except for sgGRK4.

